# Angiopoietin-like protein 2 mediates vasculopathy driven fibrogenesis in a mouse model of systemic sclerosis

**DOI:** 10.1101/2023.10.16.562467

**Authors:** Dyuti Saha, Sujaya Thannimangalath, Neha Pincha, Ravi Kiran, Akshay Hegde, Sunny Kataria, Binita Dam, Ankita Hiwale, Venkatesh Ravula, Lekshmi Minikumari Rahulan, Neha Nigam, Neha Singh, Vikas Agarwal, Praveen K. Vemula, Colin Jamora

**Affiliations:** IFOM-inStem Joint Research Laboratory, Institute for Stem Cell Science and Regenerative Medicine (DBT-inStem), Bangalore, India; Department of Biology, Manipal Academy of Higher Education, Manipal, India; Laboratory of Chemical Biology and Translational Research, Institute for Stem Cell Science and Regenerative Medicine (DBT-inStem), Bangalore, India; Department of Clinical Immunology and Rheumatology, Sanjay Gandhi Postgraduate Institute of Medical Sciences, Lucknow, India; Department of Pathology, Sanjay Gandhi Postgraduate Institute of Medical Sciences, Lucknow, India

## Abstract

Vasculopathy is a common hallmark of various fibrotic disorders including systemic sclerosis (SSc), yet its underlying etiology and contribution to fibrogenesis remain ill-defined. In SSc the vasculopathy typically precedes the onset of fibrosis and we observed that this phenomenon is recapitulated in the Snail transgenic mouse model of SSc. The vascular anomalies manifest as deformed vessels, endothelial cell dysfunction and vascular leakage. Our investigation into the underlying mechanism of this phenotype revealed that Angiopoietin-like protein 2 (Angptl2), secreted by the Snail transgenic keratinocytes, is a principal driver of fibrotic vasculopathy. In endothelial cells, Angptl2 upregulates pro-fibrotic genes, downregulates the tight junction protein Claudin 5, and prompts the acquisition of mesenchymal characteristics. Inhibiting endothelial cell junctional instability and consequently vascular leakage with a synthetic analog of the microbial metabolite Urolithin A (UAS03) effectively mitigated the vasculopathy and inhibited fibrogenesis. Thus, Angptl2 emerges as a promising early biomarker of the disease and inhibiting the vasculopathy inducing effects of this protein with agents such as UAS03 presents an appealing therapeutic avenue to reduce disease severity. These insights hold the potential to revolutionize the approach to the treatment of fibrotic diseases by targeting the vascular defects.

## INTRODUCTION

Fibrosis is a pathological condition wherein deposition of extracellular matrix components by activated fibroblasts leads to tissue stiffness and loss of function. Fibrotic diseases lead to ∼40% of deaths worldwide^1^ however the lack of effective treatments emphasizes the need for a thorough molecular understanding of fibrogenesis. In this context, the role of soluble factors such as TGFβ^2^ and immune cells and inflammation^3^ has been extensively studied for their contribution towards fibrosis development. However, targeting these pathways in clinical trials(s) have had limited success^4^, suggesting the existence of compensatory/overlapping signaling cascades that require elucidation to facilitate effective therapy development.

Fibrotic skin diseases range from localized fibrotic lesions on the skin (seen in localized scleroderma) to conditions with widespread involvement of multiple organs in systemic sclerosis (SSc)^5,6^. Like many other fibrotic tissues, a prominent hallmark of SSc are defects in blood vessels in the skin^7^. Interestingly, patient data reports that these vasculopathies manifest quite early in the disease progression^8^ suggesting their potential role in promoting fibrogenesis. Examination of patient samples from SSc has revealed that the disease is marked by early endothelial damage, followed by the accumulation of immune cell infiltrates near the affected vasculature and eventually foci of ECM deposits by activated fibroblasts^9,10^. Given the early manifestation of vascular defects in the course of fibrogenesis suggests that pathways and factors involved in mediating vascular abnormalities could potentially be used both as early biomarkers of the disease as well as therapeutic targets.

While there have been a few mouse models developed to understand the mechanistic basis of SSc progression, most do not recapitulate all the facets of the disease condition^11,12^. For example, the genetic mouse model of SSc, Tsk2/+ exhibits dermal fibrosis and inflammation but shows an absence of vascular defects^13,14^ whereas the Tsk1/+ mouse exhibits abnormal vascular tone but lacks a robust inflammatory response ^15,16^. On the other hand, chemically-induced models such as the Bleomycin induced fibrotic model^17,18^ faithfully recapitulate the dermal fibrosis and inflammation associated with SSc while only partially recapitulating the vasculature defects. Therefore, there was a need for a mouse model system that recapitulates a larger spectrum of SSc characteristics in order to elucidate the molecular mechanism of SSc vasculopathy.

We have previously developed a transgenic mouse model mimicking SSc skin by ectopically expressing the transcription factor Snail in the basal layer of keratinocytes of the mouse skin. Snail has been reported to be upregulated in various fibrotic conditions^19,20,21,22^ including SSc^23^. The Snail tg mouse skin recapitulates various histological and molecular features of SSc such as the increased dermal thickness, ECM deposition and increased inflammation. Similar to the progression of SSc, the fibrosis in the Snail tg mouse is long-lasting and originates in the skin and progresses to the involvement of internal organs^23^. Although defects related to the blood vessels such as Raynaud’s phenomenon was observed in the Snail tg skin^23^, it remains to be determined to what extent the Snail tg skin recapitulates SSc vasculopathy. The presence of SSc-like vascular defects in the Snail tg skin would position this mouse model as a platform to elucidate the underlying mechanisms of SSc vasculopathy and the contribution of vascular defects to fibrogenesis.

## RESULTS

### SSc-associated vasculopathy is recapitulated in the adult Snail transgenic skin

We have previously reported that the Snail transgenic (Snail tg) mouse recapitulates many of the diagnostic features of SSc^23,24,25^. We thus investigated whether the Snail tg mouse skin could recapitulate the various aspects of vasculopathy associated with SSc. We undertook a morphometric analysis of the cutaneous vasculopathy in 2-month-old adult Snail tg mice with the markers used to profile for human^26^ and mouse^18,27^ fibrotic skin. Staining of the back-skin of adult Snail tg mouse with the endothelial cell marker (PECAM-1, Platelet Endothelial Cell Adhesion Molecule) revealed that Snail tg skin vessels had increased morphological distortions in the form of altered vascular structures compared to its wild type counterpart (Figure 1A). Upon performing morphometric analysis, we observed a significant increase in vascular density (Figure 1B) and percentage of mean vessel area (Figure 1C) in the Snail tg skin compared to the wild type skin. Distorted vasculature is often an early driver of pathological neovascularization with hyperproliferation of endothelial cells^28^. Consistent with this, we observed a two-fold increase in the total number of endothelial cells and ∼ 9-fold increase in the number of Ki67 positive endothelial cells in the Snail tg skin vasculature compared to wild type skin (Figure 1D, E and F).

**Figure 1:**
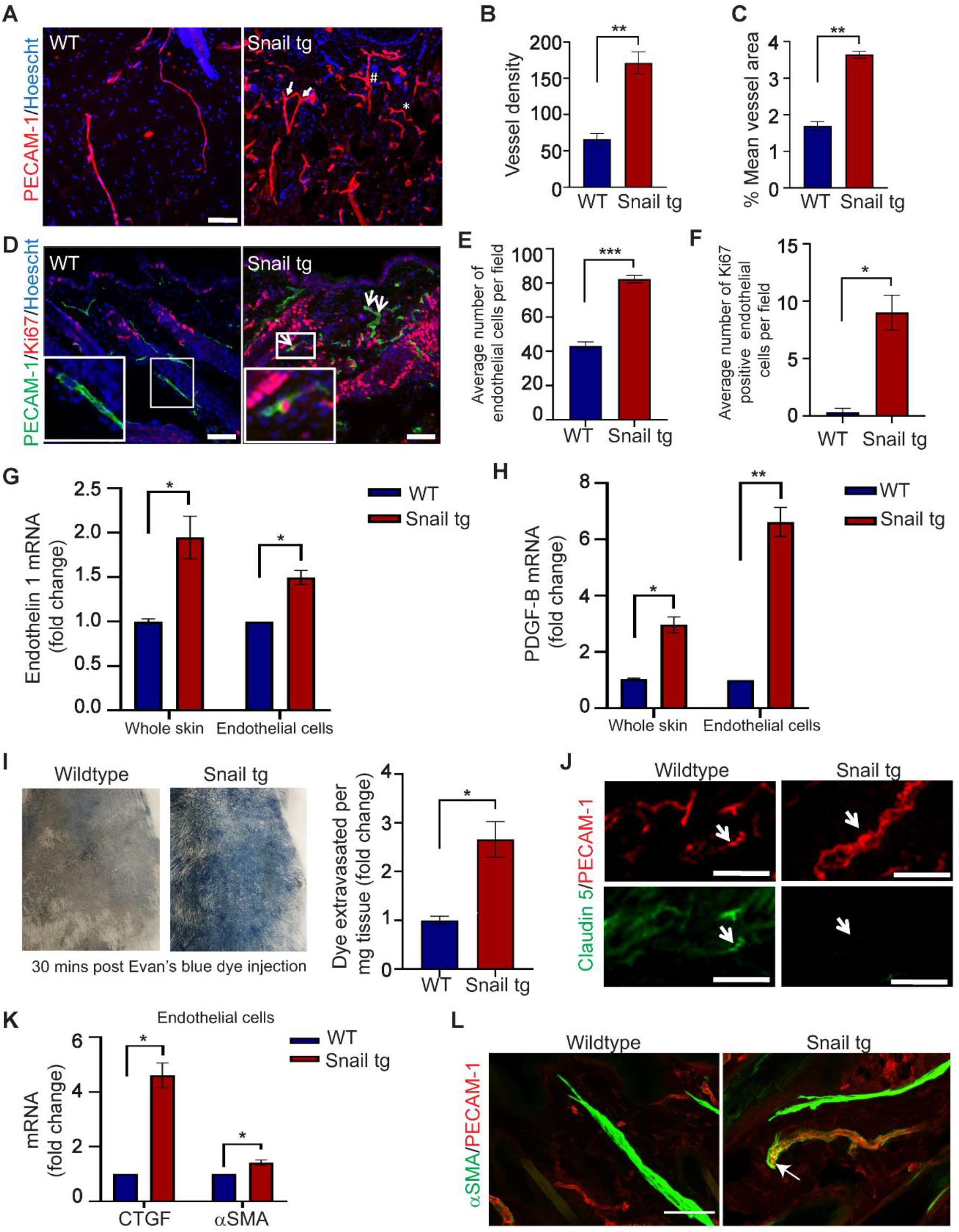
Adult Snail tg skin recapitulates vasculature abnormalities observed in scleroderma. (A) Wild type and Snail tg skin stained for PECAM-1 (red) and nuclei (blue). (B) Quantification of vessel density in wild type and Snail tg skin. (C) Quantification of mean vessel area in wild type and Snail tg skin. (D) Wild type and Snail tg skin stained for PECAM-1 (green), proliferation marker Ki67 (red) and nuclei (blue). Arrows mark PECAM-1+/Ki67+ cells. (E) Number of endothelial cells and (F) PECAM-1+/Ki67+ cells in wild type and Snail tg skin. qPCR analysis of (G) Endothelin 1 mRNA and (H) PDGF-B mRNA in whole skin and isolated dermal endothelial cells in wild type and Snail tg mice. (I) Evan’s blue dye leakage assay and quantification. (J) Wild type and Snail tg skin stained for Claudin 5 (green) and PECAM-1 (red). (K) qPCR analysis of myofibroblast markers (CTGF, αSMA) in isolated dermal endothelial cells from wild type and Snail tg mice skin. (L) Wild type and Snail tg skin stained for αSMA (green) and PECAM-1 (red). Arrow marks αSMA/PECAM-1 double positive vessel. Scale bars: 50 µm. Data are shown as mean ± SEM, p-values were calculated using unpaired Welch’s t test, * p ≤ 0.05, ** p ≤ 0.01, *** p≤ 0.001. All experiments are n=3.

We next analysed gene expression changes related to SSc vasculopathy. Endothelin-1, a marker of endothelial cell damage and dysfunction found in SSc skin^29^,^30^ is increased in the Snail tg skin (Figure 1G). Another important factor that is upregulated in scleroderma patients is Platelet-derived growth factor B (PDGF-B) that is constitutively secreted from endothelial cells and affects both perivascular cells as well as activate surrounding fibroblasts^31^. PDGF-B is likewise upregulated in the whole skin and isolated dermal endothelial cells of Snail tg mouse compared to wildtype (Figure 1H).

To investigate whether the blood vessels in Snail tg skin are functionally compromised, we examined the vascular integrity with the Evan’s blue dye leakage assay. We observed that the Snail tg skin had increased blue color indicating vascular leakage. Upon quantification, we found that Snail tg skin had a nearly three-fold increased dye extravasation into tissue from the vasculature indicating the perturbation of vessel integrity (Figure 1I). We hypothesized that disruption of endothelial cell-cell adhesion underlies the increased vascular permeability in the Snail tg skin and analysed the status of the tight junction protein Claudin 5. In the Snail tg skin, there the thickened vessels had a reduction in Claudin 5 levels (Figure 1J). Loss of endothelial barrier stability has been hypothesized to prime endothelial cells to adopt a mesenchymal phenotype^32,33^. A partial or complete acquisition of mesenchymal characteristics can cause endothelial cells to adopt features of myofibroblasts in SSc skin^34,28^. We observed upregulated mRNA expression of myofibroblast genes CTGF and αSMA in isolated endothelial cells from the Snail tg skin (Figure 1K). Also, there was presence of some αSMA/PECAM-1 double positive vessels in Snail tg skin (Figure 1L).

Overall, the Snail tg skin recapitulates the hallmark vasculature defects found in SSc thereby rendering it a useful platform to profile the developmental progression of the vasculopathy and its molecular underpinnings.

### Developmental analysis of Snail tg skin reveals that vasculopathy defects occur early in fibrogenesis

Since vasculopathy is an early phenomenon in scleroderma disease progression^7^, we hypothesized that it may have an active role in promoting fibrogenesis. To interrogate whether the Snail tg skin also exhibits early vasculopathy that precedes the thickening of the dermis, we performed a developmental analysis of the vasculopathy phenotype. Previously, we have observed increased dermal thickening (a readout for fibrosis) as early as postnatal day 9 in the Snail tg skin which increases progressively to adulthood^23^. We therefore analysed the vascular defects via PECAM-1 staining at postnatal (P) days 3, 7, and 9 and observed an increased vessel density as early as P3 in the Snail tg skin (Figure 2A). Moreover, an increase in Ki67^+^/PECAM-1^+^ double positive cells at P3 indicates the hyperproliferation of endothelial cells in the Snail tg skin (Figure 2B). Likewise, markers of endothelial damage such as Endothelin 1 (Figure 2C) and PDGF-B (Figure 2D) were upregulated in the Snail tg skin starting from postnatal day 3 suggesting that signalling pathways mediating vascular defects are initiated early in disease development in the Snail tg skin. We further tested the vascular integrity at these stages by performing the Evan’s blue dye leakage assay. As early as P3 the Snail tg skin exhibits increased vascular permeability (Figure 2E). Consistent with this vascular permeability, we observed a perturbation in the Claudin-5 beginning at P3 and appears as a punctate distribution at the surface of some vascular structures (Figure 2F). Further, the presence of some αSMA/PECAM-1 double positive vessels was observed beginning at P7 in Snail tg skin (Figure 2G)

**Figure 2:**
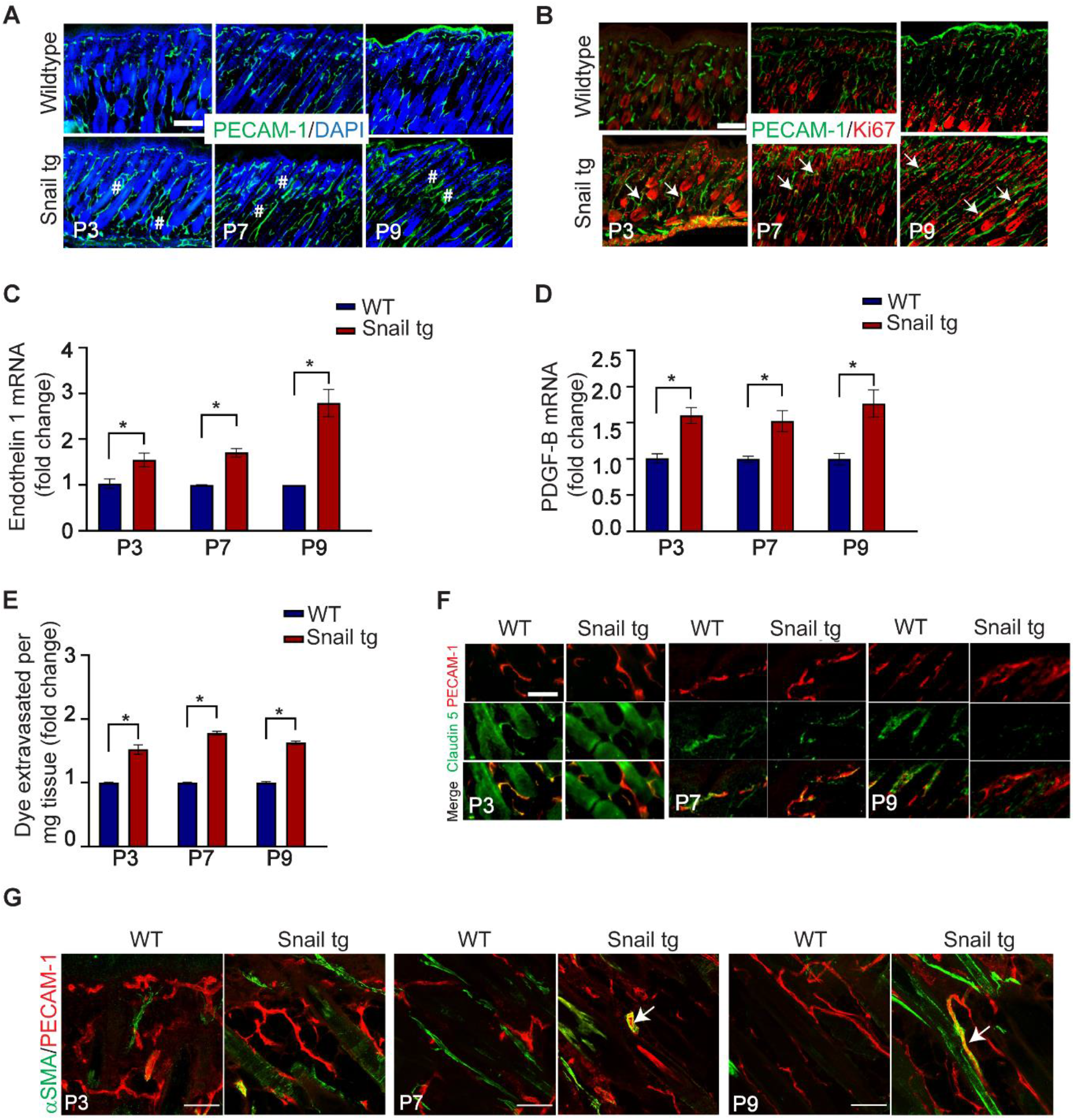
Vasculopathy phenotype occurs early in fibrogenesis. Skin from postnatal day 3, 7, and 9 wild type and Snail tg mice were stained for (A) PECAM-1 (green) and nuclei (blue). Dilated regions (#) are marked; and (B) PECAM-1 (green) and Ki67 (red). Arrows mark PECAM-1+/Ki67+ cells. qPCR analysis of (C) Endothelin 1 mRNA and (D) PDGF-B mRNA in whole skin in wild type and Snail tg mice. (E) Quantification of Evan’s blue dye leakage assay. Wild type and Snail tg skin stained for (F) Claudin 5 (green) and PECAM-1 (red), and (G) αSMA (green) and PECAM-1. Arrow marks αSMA/PECAM-1 double positive vessel. Scale bar: 100µm for (A) and (B); 50µm for (F) and (G). Data are shown as mean ± SEM, p-values were calculated using unpaired Welch’s t test, * p ≤ 0.05. All experiments are n=3.

Cumulatively our data indicate that the vasculopathy occurs in the neonatal Snail tg mouse prior to the development of significant dermal thickening, and thus recapitulates the early vascular defects that are observed in the skin of SSc patients.

### Angiopoietin-like 2 (Angptl2) secreted by Snail tg keratinocytes is necessary to induce dermal vasculopathy

Previously we have reported that extracellular factors such as PAI-1^24^ and Mindin^23^ are secreted from Snail tg keratinocytes and elicit fibrotic responses by activating dermal fibroblasts. We thus investigated whether the secretome of Snail tg keratinocytes can elicit the vascular defects found in the transgenic mouse. Conditioned media from Snail tg keratinocytes, was able to induce vascular thickening and increased density in wild type skin explants that recapitulated the vasculopathy found in the Snail tg mouse (Supplementary Figure 1A). The next question was the identity of the vasculopathy inducing factor(s) in the secretome of the Snail tg keratinocytes. Interestingly, Mindin or PAI-1 is dispensable in the vasculopathy of the Snail tg skin (Supplementary Figure 1B-D). However, we found another factor, Angiopoietin-like 2 (Angptl2), that is highly expressed (Figure 3A) and secreted (Figure 3B) from the Snail tg keratinocytes/skin at P3. Angptl2 has been reported to be involved in pathological angiogenesis in diseases such as cancer^35^. Interestingly, we found that Angptl2 is also secreted from SSc patient skin suggesting a possible role in the human disease (Figure 3C). In addition, bioinformatic analysis of Angptl2 expression in published datasets of keloid^36^ and SSc^37^ patients and fibrotic diseases of other human organs such as the lung^38^, kidney^39^ and liver^40^ revealed that Angptl2 is upregulated in the diseased skin compared to healthy skin controls (Figure 3D, Supplementary Figure 2). We next assessed whether the presence of Angptl2 was required for vasculopathy and fibrogenesis in the Snail tg adult mouse. We found that the Snail tg Angptl2 KO mouse had a reduction in vascular density (Figure 3E) and exhibited a significant decrease in vascular leakage compared to Snail tg mice (Figure 3F). Furthermore, the increased Collagen I levels and dermal thickness (readout of fibrosis) in the Snail tg skin was reduced in the absence of Angptl2 (Figure 3G and H). Overall, this data implicates Angptl2 as an important component that drives the vasculopathy and fibrotic phenotypes in the Snail tg skin.

**Figure 3:**
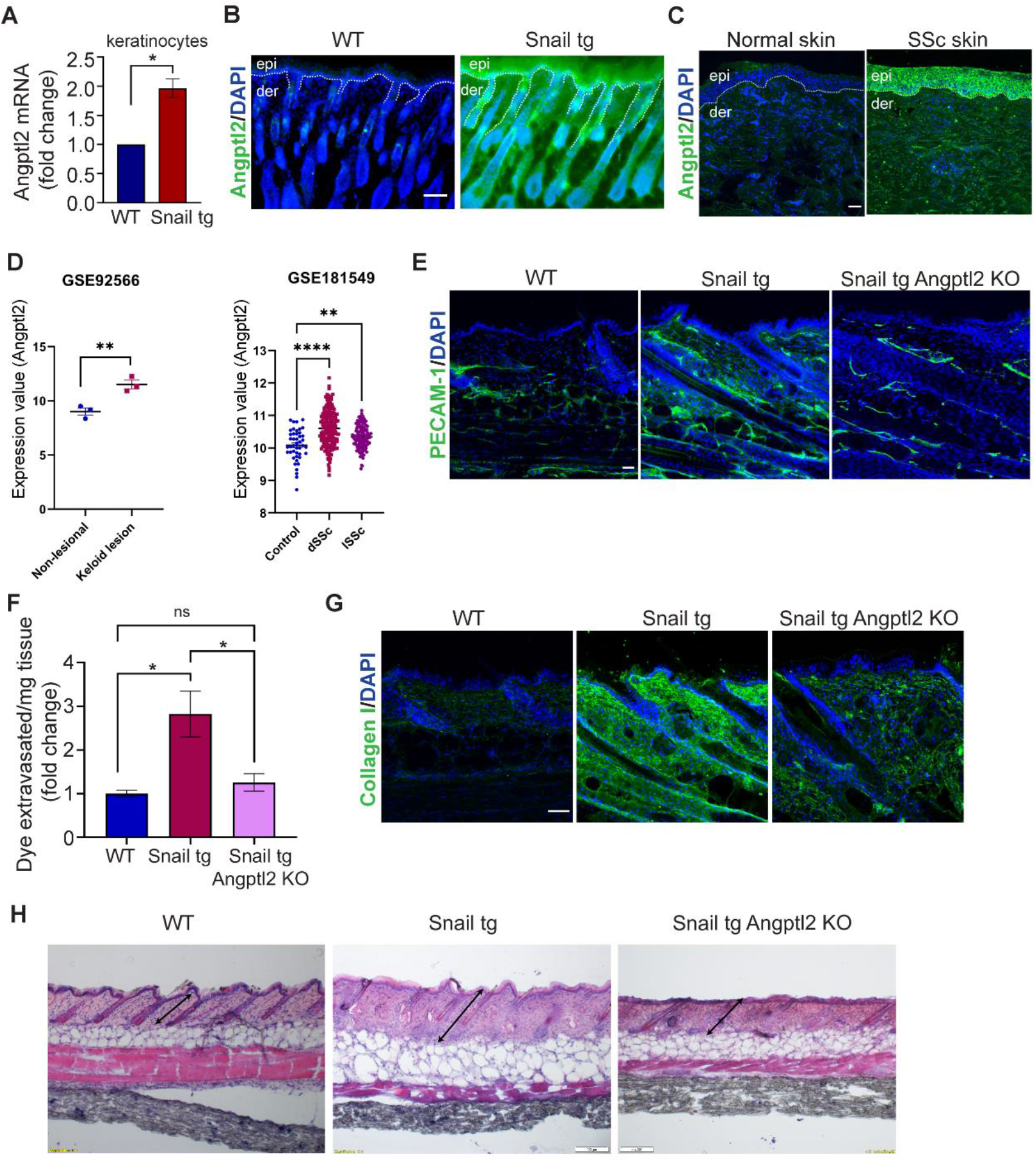
Angptl2 is necessary for fibrotic vasculopathy. (A) qPCR of Angptl2 mRNA in WT and Snail tg keratinocytes. (B) Secreted Angptl2 (in green) in WT and Snail tg skin at P3. Nuclei are marked in blue. The dotted line denotes the basement membrane separating the epidermis (epi) from the underlying dermis (der). (C) Secreted Angptl2 (in green) in normal and systemic sclerosis (SSc) skin. Nuclei are marked in blue. (D) Expression of Angptl2 in keloid lesion skin (n=3) compared to non-lesional skin (n=3) (from GSE92566) and in healthy (n=44) compared to affected forearm skin from diffuse (n=180) and localized SSc (n=115) patients (from GSE181549). The expression values are fetched from the GEO2R algorithm’s output. (E) PECAM-1 (green) staining, (F) Quantification of Evan’s blue dye leakage assay, (G) Staining for Collagen I (green) and (H) Quantification of dermal thickness in WT, Snail tg and Snail tg Angptl2 KO mouse at P60. Scale bar: 50 µm for (B), (C), (E), (G) and 100 µm for (H). Data are shown as mean ± SEM, p-values were calculated using Welch’s t test. * p ≤ 0.05, ** p ≤ 0.01. All experiments except (D) are n=3.

### Angptl2 is sufficient to drive various features of dermal vasculopathy in endothelial cells

We investigated the extent to which Angptl2 is sufficient to drive the vasculopathy observed in the skin of the Snail tg mouse. Using an ex-vivo explant assay, we found that Angptl2 can promote the dilation and swelling of vessels (Figure 4A), which recapitulates the changes in the vasculature in the Snail tg skin. Angptl2 also caused significant upregulation in mRNA levels of Endothelin 1 (Figure 4B) and PDGF-B (Figure 4C) in skin explants and the mouse endothelial cell line SVEC4-10, consistent with a role in various signalling pathways involved in the vasculopathy in the Snail tg skin. However, Angptl2 did not have a direct effect on endothelial cell proliferation (Supplementary Figure 3).

**Figure 4:**
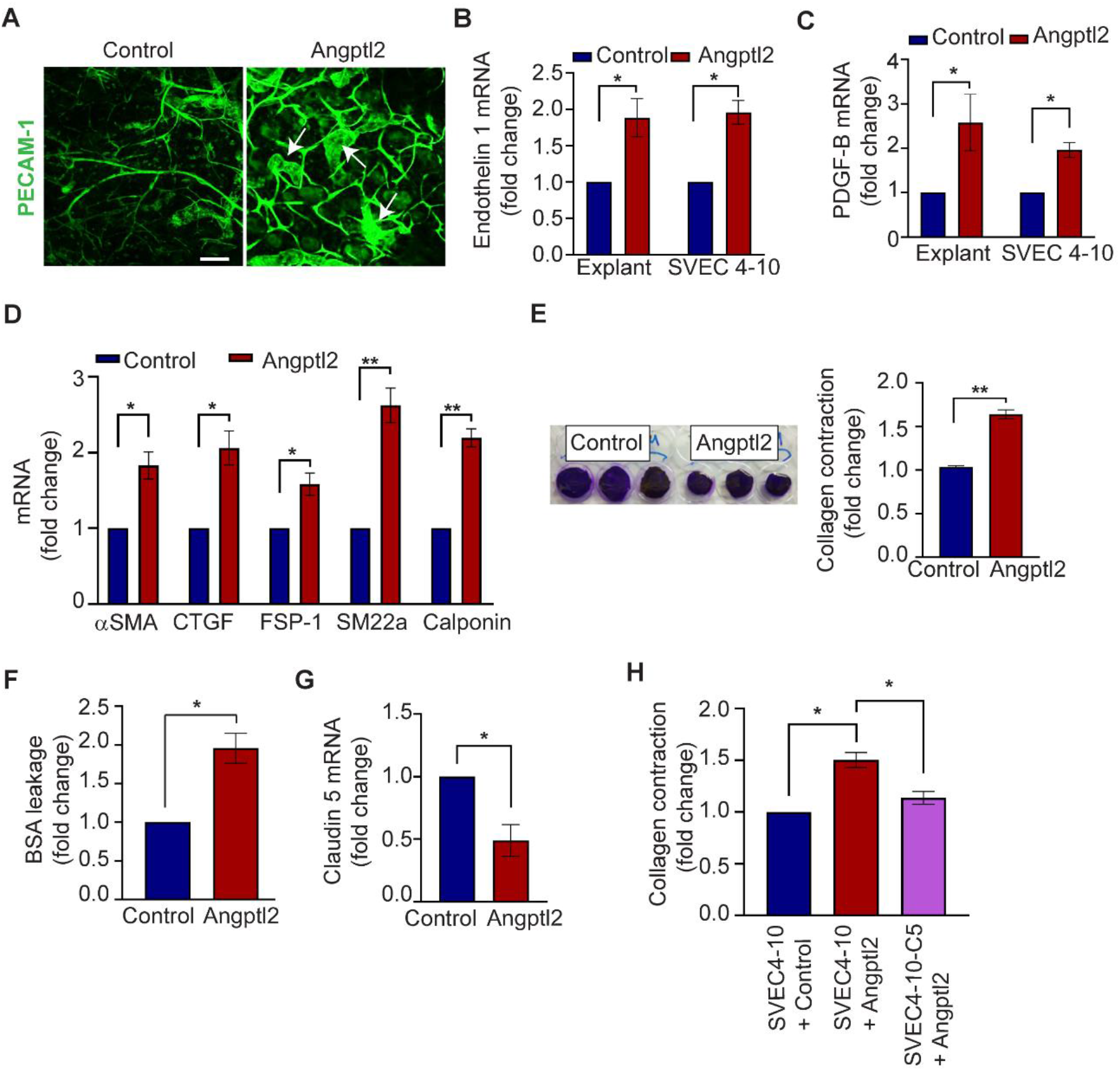
Angptl2 drives vasculopathy. (A) PECAM-1 (green) in control and Angptl2 treated skin explants. Arrows mark dilated regions. (B) Endothelin 1 and (C) PDGF-B in control and Angptl2 treated SVEC 4-10. (D) qPCR analysis of mRNA levels of myofibroblasts markers (αSMA, CTGF, FSP-1. SM22a, Calponin) in control and Angptl2 treated SVEC 4-10. (E) Collagen contraction assay (left panel) and quantification (right panel) for control and Angptl2 treated SVEC 4-10. (F) Quantification of in-vitro vascular permeability assay. (G) qPCR analysis of mRNA levels of Claudin 5 and in control and Angptl2 treated SVEC 4-10. (H) Quantification of collagen contraction assay in SVEC4-10 + Control, SVEC4-10 + Angptl2, and Claudin 5 overexpressed SVEC4-10 (SVEC 4-10-C5) + Angptl2. Scale bar: 50 µm. Data are shown as mean ± SEM, p-values were calculated using paired student t-test for (B) – (H). * p ≤ 0.05, ** p ≤ 0.01. All experiments are n=3.

Endothelial cells isolated from the Snail tg skin exhibited upregulation of myofibroblast markers, underlying our hypothesis that Angptl2 could play a role in promoting the acquisition of mesenchymal markers. Levels of myofibroblast markers (αSMA, CTGF, FSP-1, SM22α and Calponin) were upregulated in SVEC4-10 cells upon Angptl2 treatment (Figure 4D and Supplementary Figure 3). These observations are consistent with analysis of human SSc skin demonstrating the colocalization of fibroblast and endothelial markers^41^. We further examined whether these changes in the characteristic of endothelial cells resulted in alterations in vasculature functions. Angptl2 also imparted contractile behaviour in endothelial cells in collagen matrices (Figure 4E). Moreover, Angptl2 was sufficient to induce vascular leakage in a sheet of SVEC4-10 cells *in vitro* (Figure 4F).

Vascular dysfunction is often the consequence of perturbed tight junctions that is frequently due to the loss of claudin-5 expression^42^. Consistent with this, qPCR analysis revealed that mRNA levels of the tight junction protein Claudin 5 was significantly downregulated in SVEC4-10 cells treated with Angptl2 (Figure 4G). We further tested if the acquisition of mesenchymal behaviours was likewise dependent on claudin-5. To counteract the Angptl2 mediated loss of Claudin 5, a stable SVEC4-10 cell line with constitutive Claudin 5 overexpression (SVEC 4-10-C5) was generated. SVEC 4-10-C5 cells were refractory to increased contractility in collagen matrices upon Angptl2 treatment when compared to the parental SVEC 4-10 cells (Figure 4H).

Altogether, our observations suggest that Angptl2 is sufficient to drive various features of the vasculopathy observed in the Snail tg skin in-vivo. Furthermore, Claudin 5 appears to be a critical downstream component of the Angptl2-mediated vasculopathy.

### Angptl2 mediated vasculopathy in endothelial cells is abrogated by UAS03

We investigated whether upregulation of tight junction proteins such as claudin 5 through non-genetic means would be an effective mechanism to inhibit the Angptl2 driven vasculopathy. We used a synthetic analog of a gut metabolite Urolithin A (UAS03), which was previously shown to restore the disrupted epithelial barrier by upregulating tight junction proteins in a mouse model of inflammatory bowel disease^43^. However, whether this function is conserved for endothelial cell junctions was unknown. We found that UAS03 treatment reverses the decrease of claudin-5 mRNA induced by Angptl2 in SVEC 4-10 cells (Figure 5A). In line with this effect on gene expression, UAS03 could also block the Angptl2-induced vascular leakage through an endothelial sheet in vitro (Figure 5B). We further interrogated if UAS03 could counteract the other vasculopathy defects mediated by Angptl2. Increased mRNA levels of Endothelin 1 (Figure 5C) and PDGF-B (Figure 5D) induced by Angptl2 treatment were restored to control levels in the presence of UAS03 in SVEC4-10 cells. We next tested if the acquisition of mesenchymal markers associated with junctional perturbation is also inhibited upon UAS03 treatment. UAS03 treatment decreased mRNA levels of the myofibroblast markers αSMA, CTGF and FSP-1 induced by Angptl2 (Figure 5E). Consistent with the gene expression changes, UAS03 also blocked Angptl2 mediated increased contractility of endothelial cells in collagen matrices (Figure 5F). In sum, our data reveals that utilization of UAS03 is effective method in counteracting the vasculopathy induced by Angptl2 in vitro.

**Figure 5:**
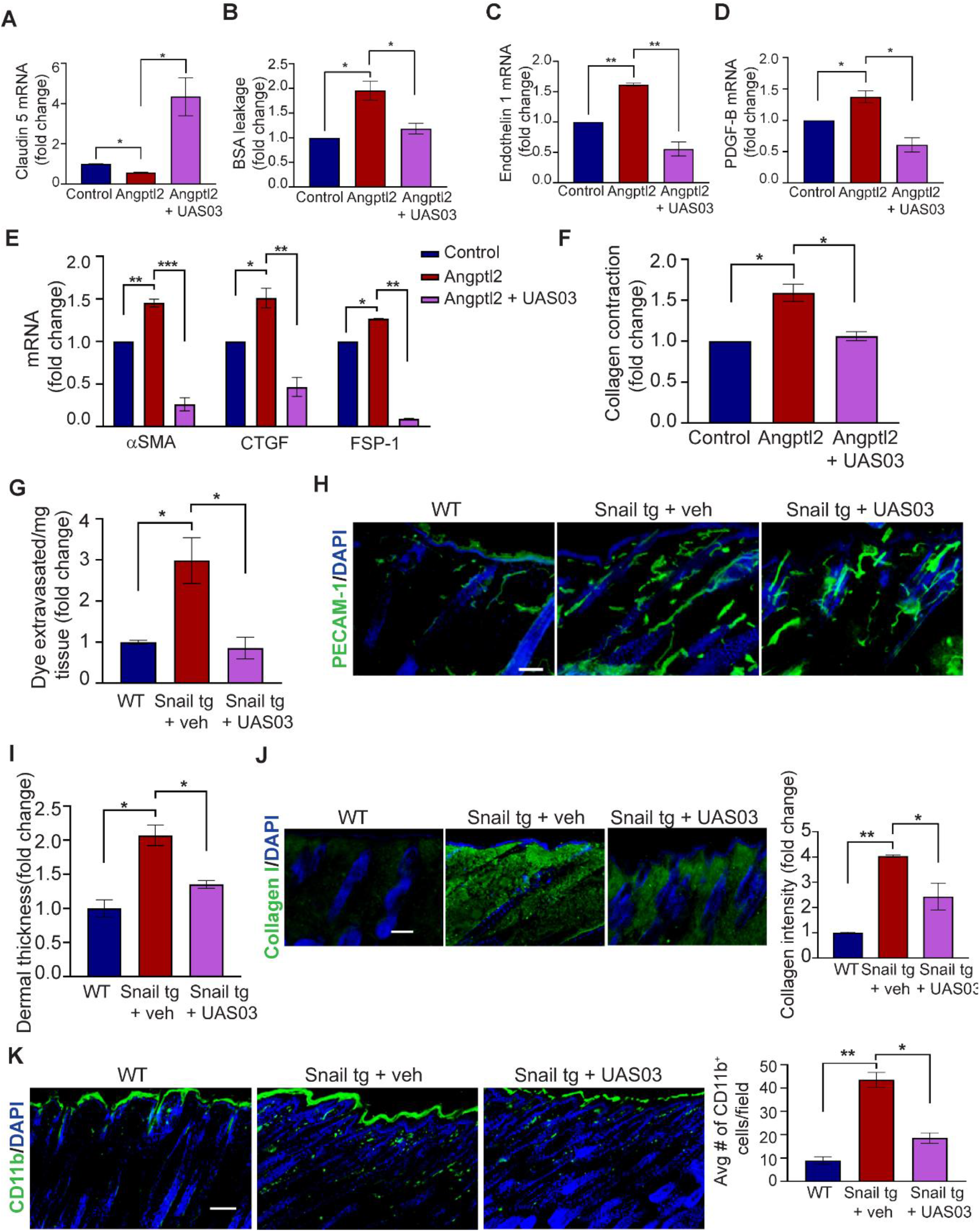
UAS03 abrogates effects of Angptl2 on endothelial cells and counteracts fibrosis. SVEC 4-10 cells were treated with control, Angptl2, or Angptl2 + UASO3 and processed for qPCR of Claudin 5 (A); in vitro vascular permeability (B); qPCR of Endothelin 1 (C), qPCR of PDGF-B (D); expression of myofibroblast markers (αSMA, CTGF, FSP-1) (E) and collagen contraction activity (F). (G) Quantification of Evan’s blue dye leakage assay. (H) Staining for PECAM-1 (green) and nuclei (DAPI) at P60 in Wild type (WT), Snail tg + vehicle control (veh) and Snail tg UAS03 injected mice. (I) Quantification of dermal thickness at P60 in WT, Snail tg + veh and Snail tg + UAS03. (J) Staining for Collagen I at P60 in WT, Snail tg + veh and Snail tg + UAS03. (K) Staining for CD11b+ cells (green) and nuclei (blue) and quantification at P9 in WT, Snail tg + veh and Snail tg + UAS03. Scale bar: 100 µm for (H) and (I), 50 µm for (K). Control and Angptl2 treatment data for (B) is taken from Figure 4(F). Data are shown as mean ± SEM, p-values were calculated using using paired Student’s t test for (A) - (F) and unpaired Welch’s t test for (G), (I)-(K), * p≤ 0.05, ** p ≤ 0.01, *** p ≤ 0.001. All experiments are n=3.

Whether UAS03 is effective in reducing the fibrotic phenotype in vivo was investigated. Intraperitoneal injections of UAS03 of mice from neonates (P3) to adulthood (P60) reduced the vascular leakage in the adult Snail tg animals to wild type levels (Figure 5G). Interestingly, vascular density remained unchanged in the UAS03 injected Snail tg skin compared to vehicle controls, suggesting that the effect of UAS03 is on the junctional integrity of the vasculature and not on the quantity of vessels (Figure 5H). We tested if the protective effects of UAS03 on the vasculature likewise impacts fibrogenesis in the Snail tg mice. We observed a significant decrease in dermal thickness (Figure 5I) and Collagen 1 protein levels (Figure 5J) in UAS03 injected Snail tg mice compared to vehicle control at P60.

Inflammation is a hallmark of fibrosis and is facilitated by the recruitment of immune cells into the tissue through leaky vessels. Among these immune cells, macrophages are well-known to be increased in many fibrotic conditions. We observed that the ∼4-fold increase of CD11b^+^ cells (which marks macrophages) in the Snail tg skin was substantially decreased in UAS03 injected tg mice (Figure 5K). We thus assessed if the UAS03-mediated decrease in the dermal fibrosis in the Snail tg skin is secondary to its effect on macrophage reduction. Interestingly, depletion of macrophage number with the chemical clodronate (Supplementary Figure 4A) was not sufficient to reduce the increased dermal thickness (Supplementary Figure 4B) and Collagen 1 protein levels (Supplementary Figure 4C) in the Snail tg mice skin compared to vehicle control.

Altogether these observations support the notion that vascular defects have an important contribution to fibrogenesis in the Snail tg mouse. Moreover, these findings position the synthetic metabolite UAS03 as a novel therapeutic strategy which can decrease vascular leakage in fibrotic skin conditions such as SSc, and thereby significantly reduce tissue fibrosis.

## DISCUSSION

Using a mouse model that recapitulates many of the clinical features of SSc^23^, we have elucidated the mechanisms underlying vasculopathy that is a hallmark of this and many fibrotic diseases. Our work positions Angptl2 secreted from Snail tg keratinocytes as a major driver of the vasculopathy observed in the fibrotic skin. Angptl2 upregulates fibrogenic molecules in endothelial cells, downregulates canonical endothelial junction protein Claudin 5 and mediates acquisition of mesenchymal characteristics. The synthetic metabolite UAS03 counteracts the effects of Angptl2 in endothelial cells and reduces fibrosis in the Snail tg skin in part by reversing the downregulation of Claudin 5 (Figure 6). These observations are consistent with previous reports that the transgenic overexpression of Angptl2 in the skin appears to phenocopy the Snail tg mouse, and in particular the vascular defects^44^.

**Figure 6:**
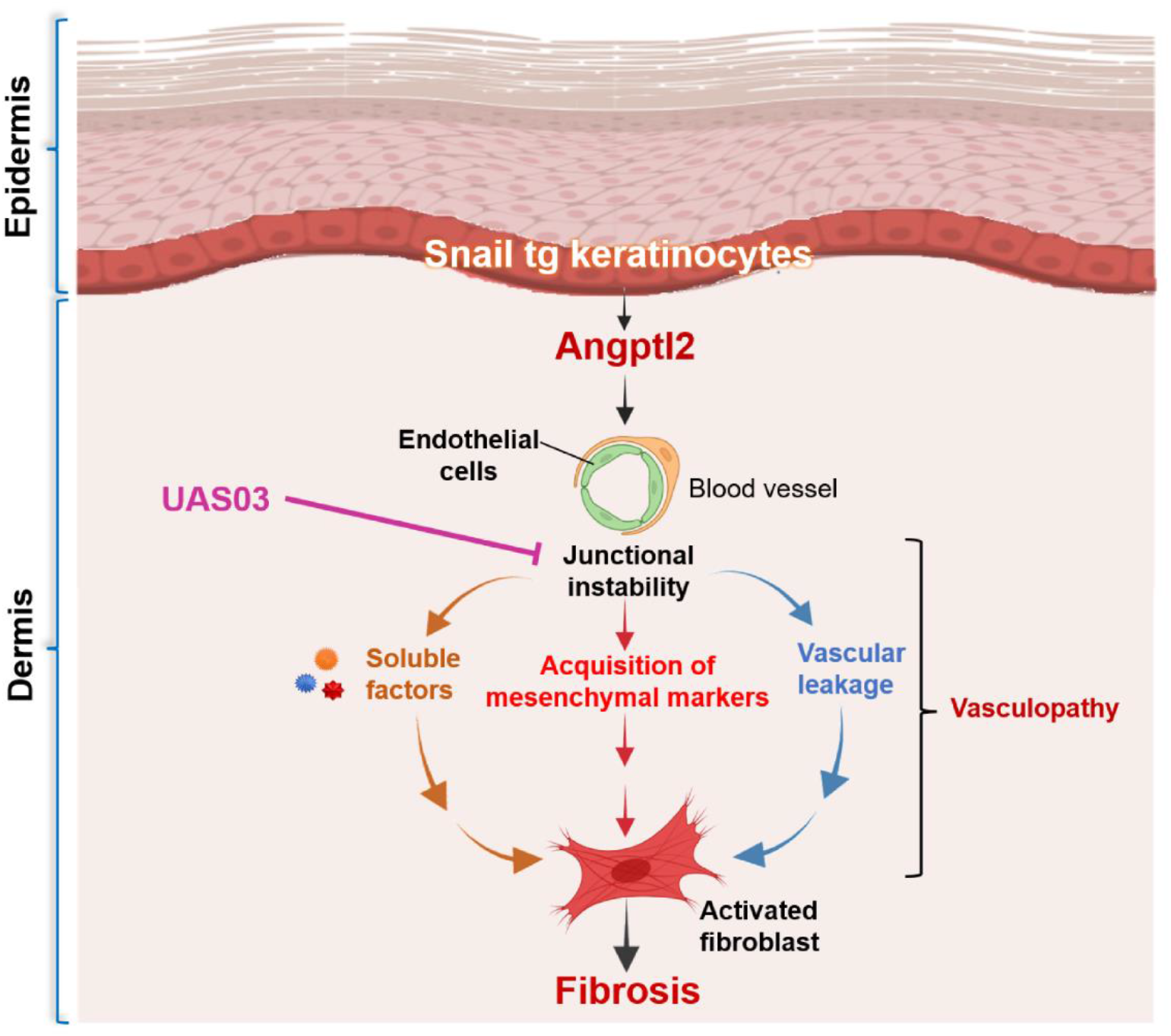
Model demonstrating vasculopathy mediated fibrosis driven by Angptl2 in Snail tg skin is counteracted by UAS03.

To date, it has not been clearly elucidated whether vasculopathy is a cause or a secondary effect of fibrosis in diseases such as SSc. Our work positions vasculopathy as an integral component driving fibrogenesis rather than a mere symptom of the disease. An outstanding question is how vasculopathy connects to other aspects of the disease pathology such as inflammation. Platelet activation has recently been hypothesized to be an important link between vasculopathy and inflammation in SSc^45^. Platelets contain granules which store various factors that can be involved in SSc vasculopathy, which includes PDGF and VEGF^46^. Intact vessel walls in unaffected vasculature usually prevents any stimuli causing platelet activation, however activated platelets can release granules containing these factors as well as molecules with pro-inflammatory activities. Endothelial dysfunction is an early event of SSc pathogenesis which leads to platelet activation^45^. Along with Angptl2-mediated junctional perturbation this can potentially be an early inducer of platelet activation setting in motion the subsequent changes in the pathogenesis of SSc.

An interesting observation in our study is that vascular defects and secretion of Angptl2 initiates in neonatal stages in the Snail tg skin. Previous reports have revealed that vasculopathy precedes other pathological features in SSc patient skin^47^ and our data concurs with this clinical observation. This leads us to propose that Angptl2 can serve as a novel biomarker for detecting SSc like pathologies at an early stage before the manifestation of dermal thickening and accumulation of ECM, which are gross indications of fibrosis. Therapeutic modalities targeting Angptl2 can also potentially be of relevance in fibrotic diseases beyond SSc. Our analysis of published datasets has revealed upregulation of Angptl2 in other fibrotic tissues such as the lung, kidney and liver (Supplementary Figure 2). In addition to fibrosis, vascular dysfunction is an integral component in the development of several pathologies such as inflammatory diseases and cancers^48,49^, which likewise exhibit elevated levels of Angptl2^50^.

Together with previous reports, our data suggests that Angptl2 does not have a role in homeostasis. Angptl2 is not required for normal vascular development and the loss of Angptl2 in mice^44^ or zebrafish^51^ do not affect the animal’s viability and fertility. Furthermore, we observed no significant effect on wound healing closure in the Angptl2 KO mouse skin (Supplementary Figure 5). This suggests that the role of Angptl2 is primarily in pathological scenarios. Consistent with this, previous reports have shown Angptl2 to have important roles in tumor angiogenesis in various cancers^35,52,53^. This indicates that a specific inhibitor to Angptl2 would likely have less side effects on normal tissue and thus have a high therapeutic potential.

As vasculopathy is a prominent feature of fibrotic diseases, previous therapies targeted factors such as Endothelin 1^54^ and VEGF^55^. Although promising in pre-clinical studies, they yielded limited success in clinical trials^54,55^. Therefore, it may be beneficial to target upstream mediators of these pathways that would theoretically have a broader effect on multiple downstream pathways. Our work has revealed that UAS03 targets multiple pathways downstream of Angptl2 leading to reduction in vasculopathy mediated fibrogenesis. The metabolite Urolithin A and its synthetic analog UAS03 have previously been reported to aid in gut epithelial junctional stability and in the reduction of inflammation in a mouse model of inflammatory bowel disease^43^. Although previous reports attributed its anti-inflammatory effects to an inhibition of macrophage activity, blocking activated macrophages via clodronate liposomes in our study was not sufficient to reduce the vasculopathy and fibrosis in the Snail tg skin (Supplementary Figure 4). Therefore, utilizing UAS03 as a therapy would not only affect inflammation but other important signaling pathways that drive fibrogenesis. Interestingly, UAS03 treated animals did not exhibit a defect in wound closure suggesting its effect is limited to pathological scenarios (Supplementary Figure 5). Therefore, our work provides important evidence supporting using UAS03 as a potential new therapy for vasculopathy mediated fibrosis development.

## METHODS

### Animal studies

The K14-Snail tg mice was described previously^56^. The Angptl2 KO mouse was developed in the Mouse Genome Engineering Facility at Bangalore Life Sciences Cluster according to previous reports^44^. Mice of both sexes were used for the experiments.

### Evan’s Blue Dye Injection Assay

Leakage of Evan’s blue dye was Evans Blue dye (1% w/v in PBS) (Sigma, E2129) was injected retro-orbitally in mice according to body weight (60mg/kg). Mice were euthanized and back skin pieces harvested. Air-dried tissue pieces were incubated with formamide at 55°C to extract the Evan’s blue dye from the tissue. Absorbance was measured at 620nm. Ng of Evan’s Blue extravasated per mg tissue was calculated from a standard curve.

### Immunostaining and histology

Skin tissue was embedded in OCT (Leica) for sectioning. 20-30μm-thick sections, whole skin mounts or cells on coverslips were fixed in 4% paraformaldehyde and used for staining. Haematoxylin/Eosin staining was used for observing tissue histology. The primary antibodies and dilutions used for immunofluorescence staining were: PECAM-1 (BD, Cat No: 550274, 1:150), Ki67 (Abcam, Cat No: AB16667, 1:200), Claudin 5 (Invitrogen, Cat No: 34-1600, 1:50), Angptl2 (R&D, Cat No: AF1444, 1:50), αSMA (Sigma, Cat No: A2547, 1:200), Collagen I (Abcam, Cat No: AB21286, 1:200), CD11b (DSHB, Cat No: M1/70.15.11.5.2, 1:200) and CD3 (Ebiosciences, Cat No:14-0032-85, 1:200). Alexa Fluor 488 and Alexa Fluor 561–labeled secondary antibodies (Jackson ImmunoResearch) were used at a dilution of 1:300. Nuclei were marked by DAPI.

### Image collection and analysis

Imaging was performed with an Olympus IX73 microscope or FV3000 confocal microscope. Images were analysed on the Fiji (ImageJ) software. For morphometric analysis of blood vessels in the Snail tg skin compared to WT, vessel density was calculated by counting total vessel profiles divided by area of tissue in mm^2^; Percentage mean vessel area was calculated as: (total vessel area/total tissue area) X 100. Collagen staining was quantified by integrated density measurement of the staining in the dermal region. Dermal thickness was quantified by measuring the area of the dermis in the H&E-stained sections.

### RNA extraction, cDNA synthesis and qRT-PCR

Total RNA was extracted from skin biopsies and cell lysates using RNAiso Plus Reagent (Takara, Cat No: 9109) according to the manufacturer’s protocol. cDNA was prepared using PrimeScript cDNA synthesis kit (Takara, Cat No: 2680A) according to the manufacturer’s instructions. Quantitative PCR (qPCR) was performed using PowerUp SYBR Green Master Mix (Applied Biosystems, Thermo Fisher Scientific, Cat No: A25742) in a Bio-Rad CFX384 machine. TBP expression was used for normalization. The primers used are listed in Table 2.

**Table 2.**
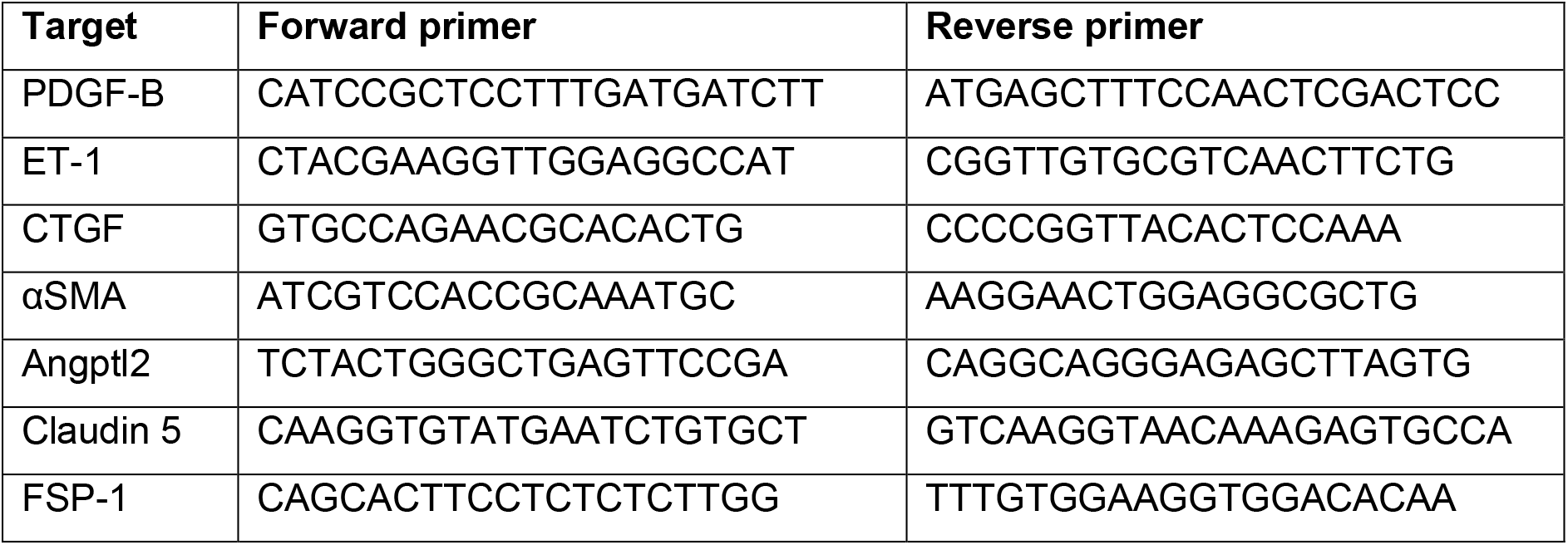
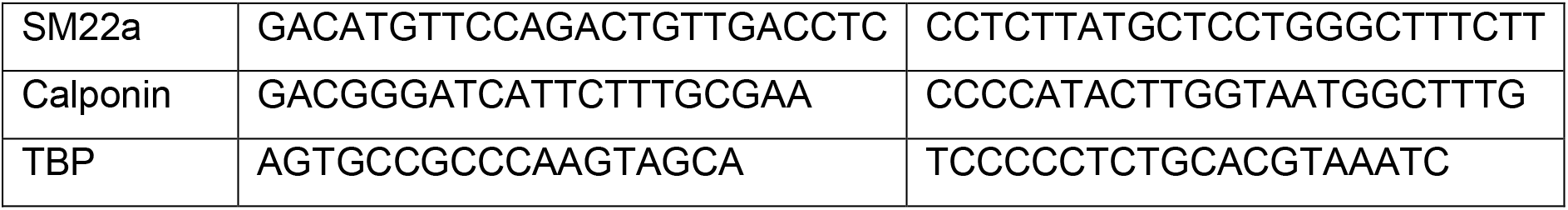
List of quantitative real time PCR primers.

### Isolation of endothelial cells from mice skin

Back skin of adult (P60) wildtype and Snail tg mice were taken, all the hair removed and chopped finely. The tissue pieces were incubated with collagenase for 2 hours at 37°C under shaking conditions and cells pelleted (350g 8 mins) after straining through a 70-micron strainer. Cell pellet was resuspended in Phosphate buffered saline (PBS) containing 1% Fetal bovine serum. The cell suspension was incubated with PE conjugated PECAM-1 antibody (MACS Miltenyi Biotech, Cat No: 130-123-675, 1:50) at 4°C for 30 mins. After a PBS wash, cells were pelleted again (350g 8 mins), resuspended in PBS and passed through a 40-micron strainer to obtain a single cell suspension. The cells were subjected to flow cytometry using FACS Aria Fusion to obtained the PECAM 1^+^ positive cells.

### Cell culture and preparation of conditioned media

Mouse endothelial cell line SVEC4-10 (ATCC® CRL-2181™) was purchased from ATCC. A Chinese hamster ovarian (CHO) cell line stably secreting Angiopoietin-like protein 2 was developed by transfecting pcDNA3.1 plasmid containing the mouse Angptl2 gene. Conditioned media from CHO cells with an empty plasmid was used as control in all the experiments. All cell lines were maintained in Dulbecco’s Modified Eagle’s Media (DMEM) supplemented with 10% Fetal bovine serum, antibiotic PenStrep, sodium pyruvate and non-essential amino acids in 5% CO2 at 37°C. For preparation of conditioned media, CHO cells secreting Angptl2 and control CHO cells were incubated with media containing 2% FBS for 3 days and the conditioned media collected and snap-frozen before storing at -80°C. For all treatments of SVEC 4-10 endothelial cells this conditioned media was used directly to treat for 48 hours. For all staining of SVEC 4-10 cells, cells were grown on a coverslip coated with 50µg/ml Collagen I solution till they formed an endothelial sheet and treatments done subsequently.

Isolation and culture of primary epidermal keratinocytes was done as described previously^57^. Cells were grown to confluency and RNA extracted as described above. For collection of conditioned media from keratinocytes to treat explants, cells were grown in low Calcium E-media as reported previously^58^. Serum free media was added to confluent plates of cells and conditioned media collected after 16 hours.

### UAS03 treatments

UAS03 formulation was obtained from Dr. Praveen Vemula at inStem. For in-vivo experiments, UAS03 (20 mg/kg body weight) was injected interperitoneally once a week starting from post-natal day 3 to 60 when mice were sacrificed. For all in-vitro experiments with SVEC 4-10 cells, 50µM UAS03 in DMSO formulation was used.

### In-vitro permeability Assay

SVEC 4-10 cells were grown on 24-well transwell inserts for 72 hours to form a confluent monolayer. 100000 cells were seeded per insert. Cells were treated with Control, Angptl2 or UAS03 for 48 hours. Both chambers were washed with PBS and plain DMEM added. 40mg/ml BSA was added in the top chamber and cells incubated at 37℃ for 2 hours. BSA concentration was measured in the bottom chamber using Bradford assay. Amount of BSA permeated across the endothelial layer was calculated from a standard curve.

### Ex-vivo treatment of skin explants and ears with Angptl2

Wild-type adult mouse ears or back skin explants were obtained immediately after sacrificing mice by cervical dislocation, placed in PBS with antibiotics and antifungal drugs (PenStrep, Gentamicin, Fungizone) for 2 hours. Explants were transferred to cell culture plates (dermis side down) containing CHO cell line stably secreting Angiopoietin-like protein 2 (ANGPTL2) or control CHO cells. After 5 days explants were removed and staining and RNA extraction done as described above.

### Analysis of published datasets for Angptl2 gene expression

The normalized expression of Angptl2 in Keloids, SSc, lung with idiopathic pulmonary fibrosis, kidney glomeruli with lupus nephropathy and liver with fibrosis post-transplant was obtained from publicly available microarray data (GSE92566, GSE181549, GSE48149, GSE32591, GSE145780). GEO2R tool (NCBI) was used to analyse the expression level difference between two groups. Welch’s t-test used to identify significance level between two groups.

### Collagen contraction assay

Assay was performed as described previously^24^. SVEC 4-10 or SVEC 4-10 cells with Claudin 5 overexpression were used. Briefly, cells were seeded at a density of 120000 cells in plugs made of Rat tail collagen I (Millipore; 08-115) in a 24-well plate and treatment done with Control, Angptl2 and/or UAS03. Collagen plugs were stained with crystal violet and contraction measured from the gel images after 72 hours. The values are represented as fold change of increase in contraction (1/area of collagen gel).

### Human samples

Scleroderma patient samples and appropriate control skin samples were obtained from Dr. Vikas Agarwal, Sanjay Gandhi Post Graduate Institute of Medical Sciences, Lucknow, India. Skin-punch biopsies were taken from the arms of patients diagnosed with diffuse systemic sclerosis or nonsystemic sclerosis.

### Wounding studies

Twelve-week-old 1) WT mice injected with control or UAS03, or 2) WT and Angptl2 KO mice were wounded with 5-mm punch biopsies, and wounded skin imaged till wound closure. Wound areas were calculated as percentage open wound.

### Statistical analysis

Comparison of 2 groups was done using either unpaired Welch’s t-test (for in-vivo experiments) or paired Students t-test (for in-vitro experiments with endothelial cells). Data are shown as mean ± SEM. GraphPad Prism 5.02 was used for all statistical analyses. P values less than 0.05 were considered significant.

### Study approval

Animal work in the Jamora lab was approved by the Institutional Animal Ethics Committee (INS-IAE-2019/06[R1]). Acquisition and processing of the human tissue were conducted according to the protocol approved by the Institutional Review Board of the Sanjay Gandhi Postgraduate Institute of Medical Sciences (Lucknow, India). Informed consent was acquired from all patients for skin sample collection and experimentation. All experimental work was done with approval of the Institutional Biosafety Committee at inStem.

## Data availability

This paper does not include any next-generation sequencing datasets and has not used or made any particular codes for large-scale data analysis.

## Author contributions

Conceptualization: DS, CJ; Methodology: DS, ST, NP, CJ; Investigation: DS, ST, NP, RK, AH, SK, BD, AH, VR; Formal Analysis: DS, ST, RK, SK; Visualization: DS; Resource: LMR, NN, NS, VA, PKV; Funding Acquisition: CJ; Supervision: PKV, CJ; Writing - Original Draft Preparation: DS, CJ; Writing - Review and Editing: DS, CJ.

## Acknowledgments

The authors would like to thank members of the Jamora laboratory and John Varga for critical review of the manuscript and Srikala Raghavan, Tina Mukherjee and Maneesha Inamdar for insightful discussions. This work was supported by core funds from inStem and grants from the Department of Biotechnology of the Government of India (BT/PR8738/AGR/36/770/2013, BT/PR32539/BRB/10/1814/2019). Animal studies were partially supported by the National Mouse Research Resource grant BT/ PR5981/MED/31/181/2012;2013-2016;2018 and 102/IFD/SAN/5003/2017-2018 from the Department of Biotechnology. We thank the staff of the Animal Care and Resource Centre and Mouse Genome Engineering Facility (Aurelie Jory, Vaishak Nair Penar, B. Sangeetha, Lalitha K S) and the Central Imaging and Flow Cytometry Facility at the Bangalore Life Science Cluster.

