## Supplementary Figures for "Angiopoietin-like protein 2 mediates vasculopathy driven fibrogenesis in a mouse model of systemic sclerosis"

### Supplementary Figure 1

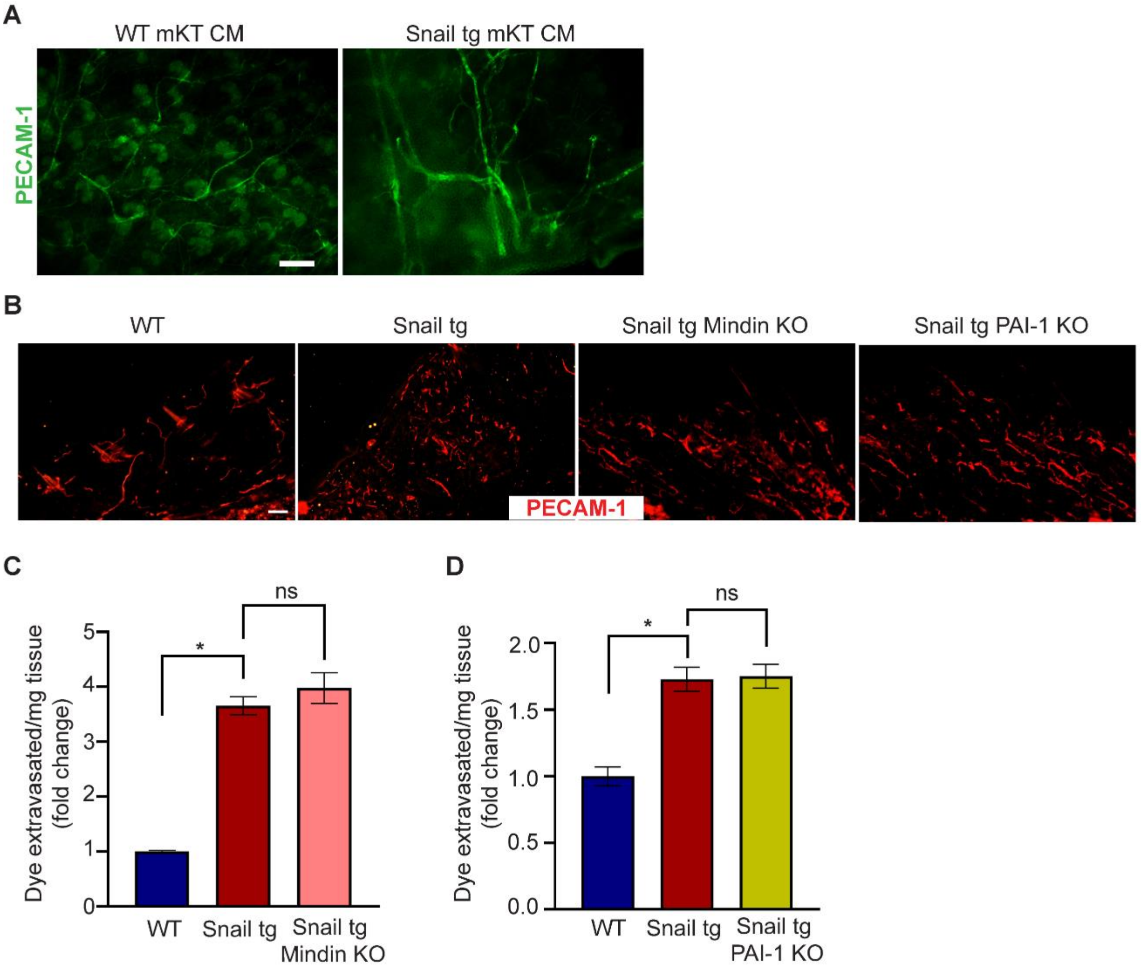


**Supplementary Figure 1: Mindin and PAI-1 do not mediate the vasculopathy in the Snail tg skin.** (A) PECAM-1 (green) in skin explants treated with WT or Snail tg mouse keratinocyte (mKT) conditioned media. Scale bar: 100 µm. (B) PECAM-1 (red) in WT, Snail tg, Snail tg Mindin KO and Snail tg PAI-1 KO skin at P60. (C), (D) Quantification of Evan's blue dye leakage assay. Scale bar: 50 µm. Data are shown as mean ± SEM, p-values were calculated using unpaired Welch's t test. * $p \leq 0.05$, ns >0.05. All experiments are n=3.

### Supplementary Figure 2

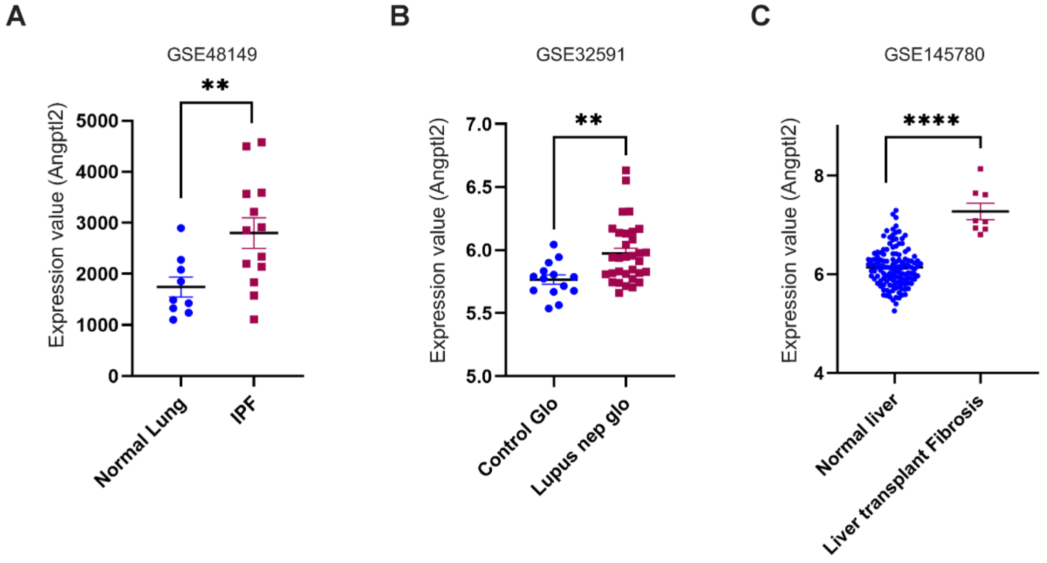


**Supplementary Figure 2: Angptl2 is upregulated in human fibrotic diseases.** Expression of Angptl2 in (A) lung with idiopathic pulmonary fibrosis (n=13) compared to normal lung tissue (n=9) (from GSE48149), (B) kidney glomeruli with lupus nephropathy (n=32) compared to control glomeruli (n=14) (from GSE32591), (C) Liver with fibrosis post-transplant (n=8) compared to normal liver tissue (n=129) (from GSE145780). The expression values are fetched from the GEO2R algorithm's output. Data are shown as mean ± SEM, p-values were calculated using unpaired Welch's t test. **** $p < 0.0001$, ** $p < 0.001$.

### Supplementary Figure 3

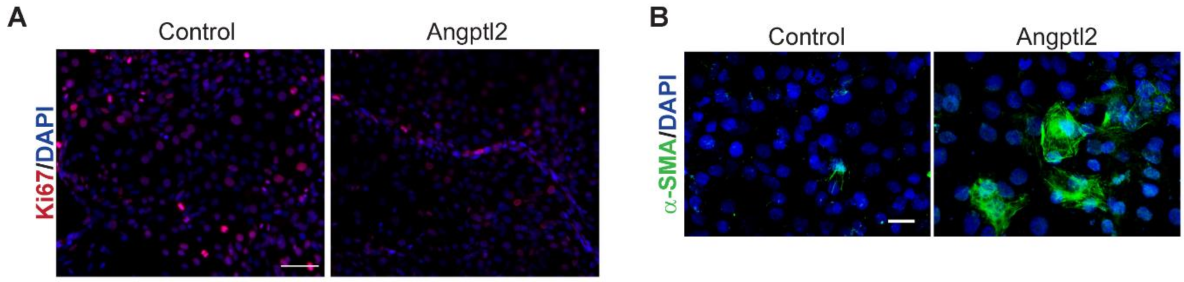


**Supplementary Figure 3: Effect of Angptl2 on endothelial cells.** (A) Ki67 (red) in control and Angptl2 treated SVEC 4-10. (B) αSMA (green) in control and Angptl2 treated SVEC 4-10 cells. Nuclei are marked in blue. Scale bar: 50 μm.

### Supplementary Figure 4

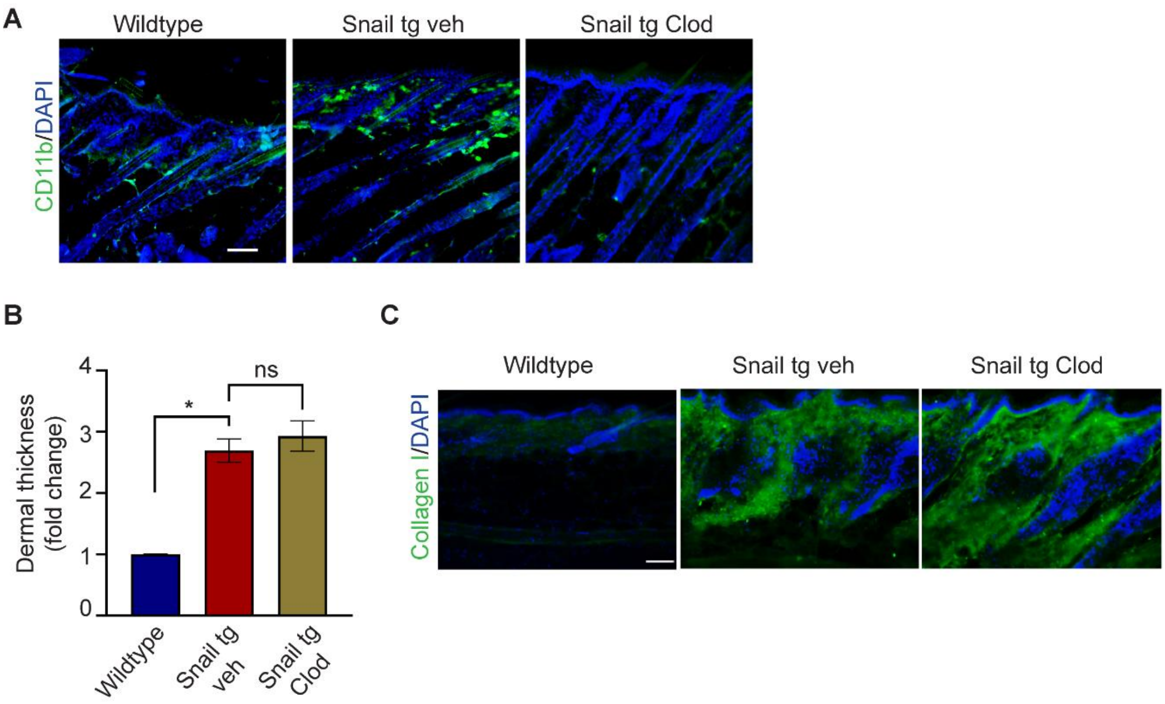


**Supplementary Figure 4: Depleting macrophages does not reduce fibrosis.** (A) Staining for macrophages marked by CD11b at P60 in wild type, Snail tg control and Snail tg Clodronate injected mice. (B) Quantification of dermal thickness at P60 in wild type, Snail tg vehicle control and Snail tg Clodronate injected mice. (C) Staining for Collagen I at P60 in wild type, Snail tg control and Snail tg Clodronate injected mice. Scale bar: 50 µm. Data are shown as mean ± SEM, p-values were calculated using unpaired Welch's t test, * $p \leq 0.05$, ns >0.05. All experiments are n=3.

### Supplementary Figure 5

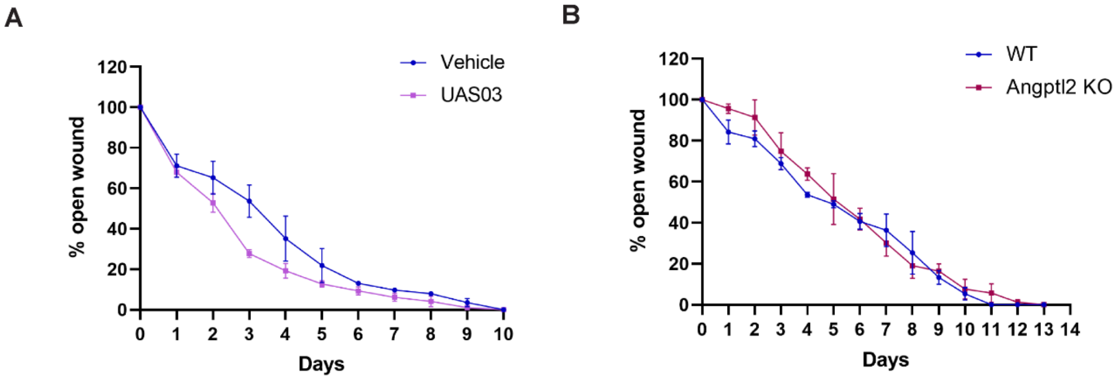


**Supplementary Figure 5:** Wound closure kinetics in (A) UAS03 treated and (B) Angptl2 KO mice compared to wildtype counterparts.
